# The secretory fate of flavivirus NS1 in mosquito cells is influenced by the caveolin binding domain

**DOI:** 10.1101/2019.12.16.879031

**Authors:** Romel Rosales Ramirez, Juan E. Ludert

**Affiliations:** Department of Infectomics and Molecular Pathogenesis, Center for Research and Advanced Studies (CINVESTAV-IPN) Mexico City, Mexico

**Keywords:** Dengue, Zika, flavivirus, NS1, caveolin, viral protein trafficking, unconventional secretion, mosquito cells

## Abstract

Flaviviruses of major medical importance worldwide such as dengue (DENV), Zika (ZIKV), and yellow fever (YFV) viruses are transmitted by mosquitoes *Aedes sp*. The non-structural protein 1 (NS1) of these flaviviruses is secreted from the infected cells using different secretion routes depending on the cell and virus nature. The NS1 of DENV and ZIKV contain in the hydrophobic region a conserved caveolin binding domain (CBD) (ΦXXΦXXXXΦ), which is not conserved in YFV NS1. To ascertain the role of the CBD in the secretory route followed by flavivirus NS1, expression vectors for the NS1 of DENV2, ZIKV and YFV were constructed. Using site-directed mutagenesis, substitutions were made in the aromatic residues within CBD; in addition, the full domain was replaced by those of other flaviviruses, creating chimeras in the CBD of NS1. Substitutions of the aromatic residues to Ala or Thr, or CBD chimeras, results in increased sensitivity of NS1 secretion to brefeldin A treatment, indicating a change to a classical secretion pathway. Likewise, the insertion of the DENV/ZIKV CBD into the recombinant *Gaussia*-Luciferase results in a loss of sensitivity to BFA treatment, in luciferase secretion. These results suggest that the CBD sequence is a molecular determinant for the unconventional secretory route followed by DENV and ZIKV NS1 in mosquito cells. However, the cellular components that recognize the CBD in the NS1 of DENV and ZIKV and redirect them to an unconventional route and if this secretion route confers unique functions to NS1 within the vector mosquito are aspects currently unknown.

**Importance:** Flaviviruses are an important cause of mosquito borne diseases to humans. We have previously demonstrated that the non-structural protein 1 from dengue and zika virus are secreted efficiently from mosquito cells using an unconventional route, that depends on caveolin and molecular chaperones. In this work, we show evidence indicating that a caveolin binding domain, well conserved and exposed in dengue and Zika virus NS1, but absent in other flaviviruses such as yellow fever virus or West Nile virus, is important in determining the unconventional secretion pathway followed by dengue and zika virus NS1 in mosquito cells. The unique secretory pathway followed by NS1 in mosquito cells may result in distinctive viral-cellular protein associations required to facilitate viral infection in the mosquito vector. To identify viral and cellular elements that could disturb the traffic of dengue and Zika virus NS1 may be important to design of strategies for vector control.

## INTRODUCTION

The *Flaviviridae* family includes many significant viral human pathogens, including yellow fever virus (YFV), dengue virus (DENV), Zika virus (ZIKV), West Nile virus (WNV), Japanese encephalitis virus (JEV), and tick-borne encephalitic virus (TBEV). The flavivirus virion particles are small (∼50 nm) and contain a single-stranded positive RNA genome of nearly 11,000 bases in length (1). DENV is the only flavivirus with four serotypes (DENV1-4) and infection with any of them can cause dengue fever or severe dengue. DENV, ZIKV and YFV are transmitted by *Aedes* mosquitoes and circulate in tropical and sub-tropical regions of the globe (2). There are several environmental, demographic and eco-logical reasons to believe that either novel or known flaviviruses will continue to emerge. In this respect, the success of vaccination against YFV has been temperate by difficulties encountered when vaccination was launched against DENV(3). In particular, the presence of four DENV serotypes has complicated vaccine design because incomplete protection against one serotype may influence the disease outcome, once infection is established by a distinct serotype, through a process referred to as antibody-mediated disease enhancement (4).

The flavivirus genome encodes only one open reading frame that is translated as one large polyprotein. The polyprotein is then cleaved by host and viral proteases to release individual viral proteins. The genome of most flavivirus encodes for three structural (C, E, prM/M) and seven non-structural (NS) (NS1, NS2A, NS2B, NS3, NS4A, NS4B, NS5) proteins (5). In DENV, the NS1 protein act as an scaffolding protein that anchors the replication complex to the ER membrane and interacts physically with NS4B (6). The NS1 protein is a 352-amino-acid polypeptide with a molecular weight of 46–55 kDa, depending on its glycosylation status. The NS1 protein exists in multiple oligomeric forms and is found in different cellular locations: a cell membrane-bound form in association with virus-induced intracellular vesicular compartments, on the cell surface and as a soluble secreted hexameric lipoparticle (6). The NS1 monomeric form rapidly dimerizes in the endoplasmic reticulum (ER), then three dimeric forms of NS1 arrange to form an hexamer (7). The hexameric form of NS1 shows an open barrel form filled with lipids and cholesterol which resemble the lipid composition of the HDL particle (8). NS1 associated pathogenesis, comprising several diverse mechanisms, in the vertebrate host has been described (9, 10). However, the possible NS1 pathogenic effects within the mosquito vector are a matter of study and are not been well understood yet (11).

In a previous work, DENV and ZIKV NS1 secretion in infected mosquito cells was associated to a caveolin-1 (CAV-1) dependent, unconventional secretory pathway that bypasses the Golgi-complex (12, 13). In contrast, YFV NS1 is secreted in both vertebrate and mosquito cells using Golgi-dependent, classical secretory pathway (13, 14). Furthermore, it was determine that NS1 secretion in mosquito cells is dependent on the caveolin chaperone complex (CCC) (13). The interaction of proteins with CAV-1, which seems to be important in the recruitment of proteins to the caveolar domains and, therefore, in the formation of microenvironments rich in interactive signaling molecules, is believed to be mediated through the interaction of an N-20 amino acid terminal region in the caveolin molecule, known as the caveolin scaffolding domain, and the CBD present in the likely caveolin binding proteins (15–17). The CBD is defined as a sequence of three or four aromatic residues separated by unspecified amino acids (ΦXΦXXXXΦ, ΦXXXXΦXXΦ or ΦXΦXXXXΦXXΦ, where Φ is any aromatic amino acid) (15). The NS1 of all 4 DENV serotypes and ZIKV present a well conserved and exposed caveolin-binding domain (FXXFXXXXW) (12), which is absent in others mosquito borne flaviviruses such as YFV, WNV and JEV. The DENV and ZIKV NS1 CDB is located in the connector subdomain, in the so-called “butter fingers”, a hydrophobic region which creates a protrusion with a hydrophobic surface, that in the dimer is in close contact with the lipid bilayer and in the hexamer is in contact with the lipids in the center (18).

The previous observation that DENV and ZIKV associate with CAV-1 and the CCC, and are secreted bypassing the Golgi-complex while the YFV NS1 do not show such associations and is secreted by a classical secretory route (12, 13), indicate that the association of DENV and ZIKV NS1 with the CCC, and the use of an unconventional secretory route, may respond to the presence of the CBD. Therefore, in this work, we examined in more detail the relationship between the CBD sequence in the flavivirus NS1 and its secretion pathway. By using site point mutations in aromatics residues and the generation of chimeras, data was obtained indicating that indeed the unconventional secretion route followed by DENV and ZIKV NS1 in mosquito cells is influenced by the CDB. This work brings new knowledge about how the secretory machinery in mosquito cells senses sequences in proteins and directs them to its secretion pathway.

## RESULTS

To determine whether the presence of CBD in the NS1 sequence of DENV or ZIKV influences the secretion path of NS1 in mosquito cells, first plasmids with the recombinant DENV, ZIKV and well as YFV, for comparison, interest were generated. In addition, to site-directed mutants, clones with full substitution of the CBD between different NS1s and NS1 chimeras with *Gau*-Luc were generated. Figure 1 shows the general experimental scheme followed by in this work and Table 1 the list of all primers used.

**Table 1.**
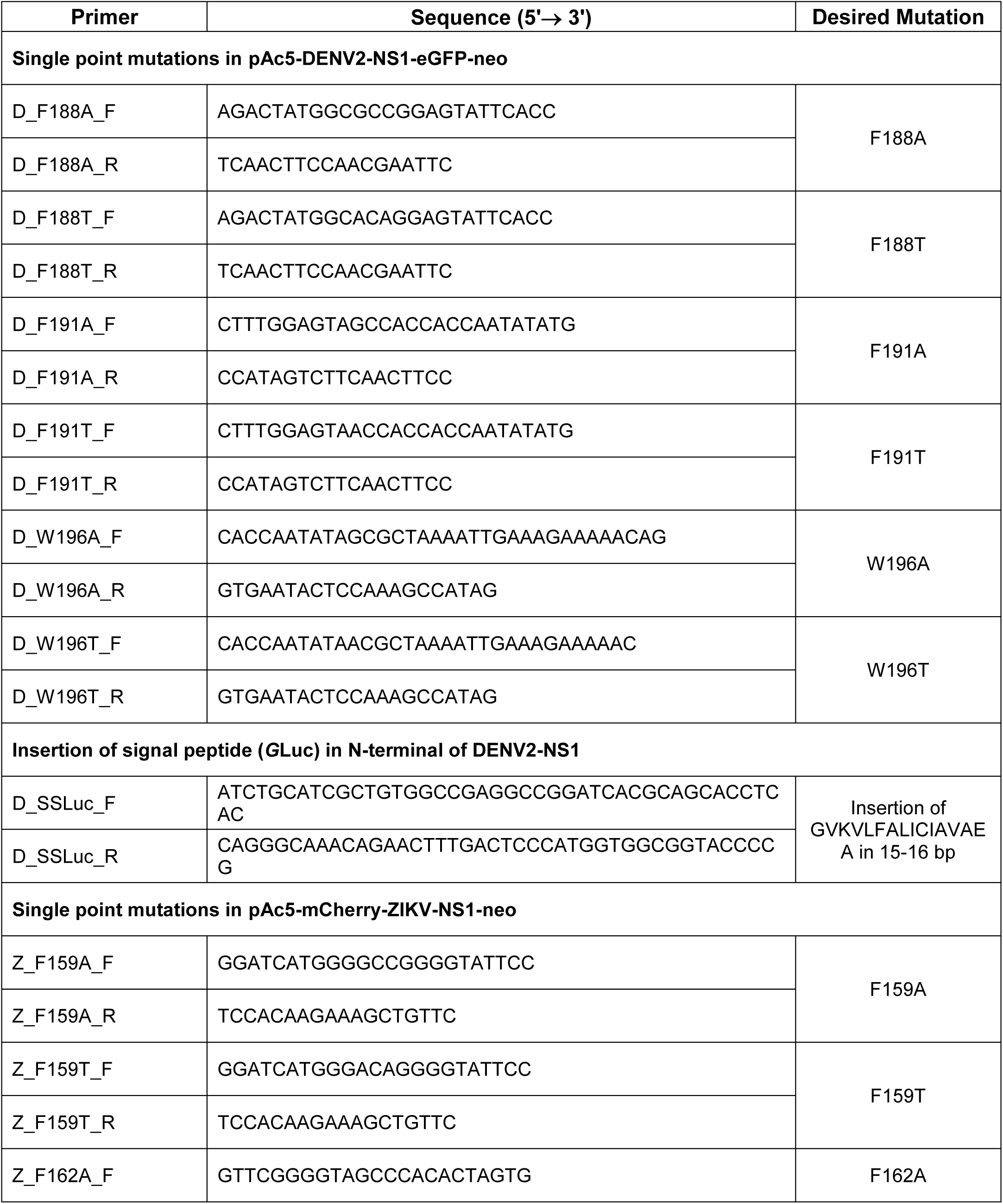
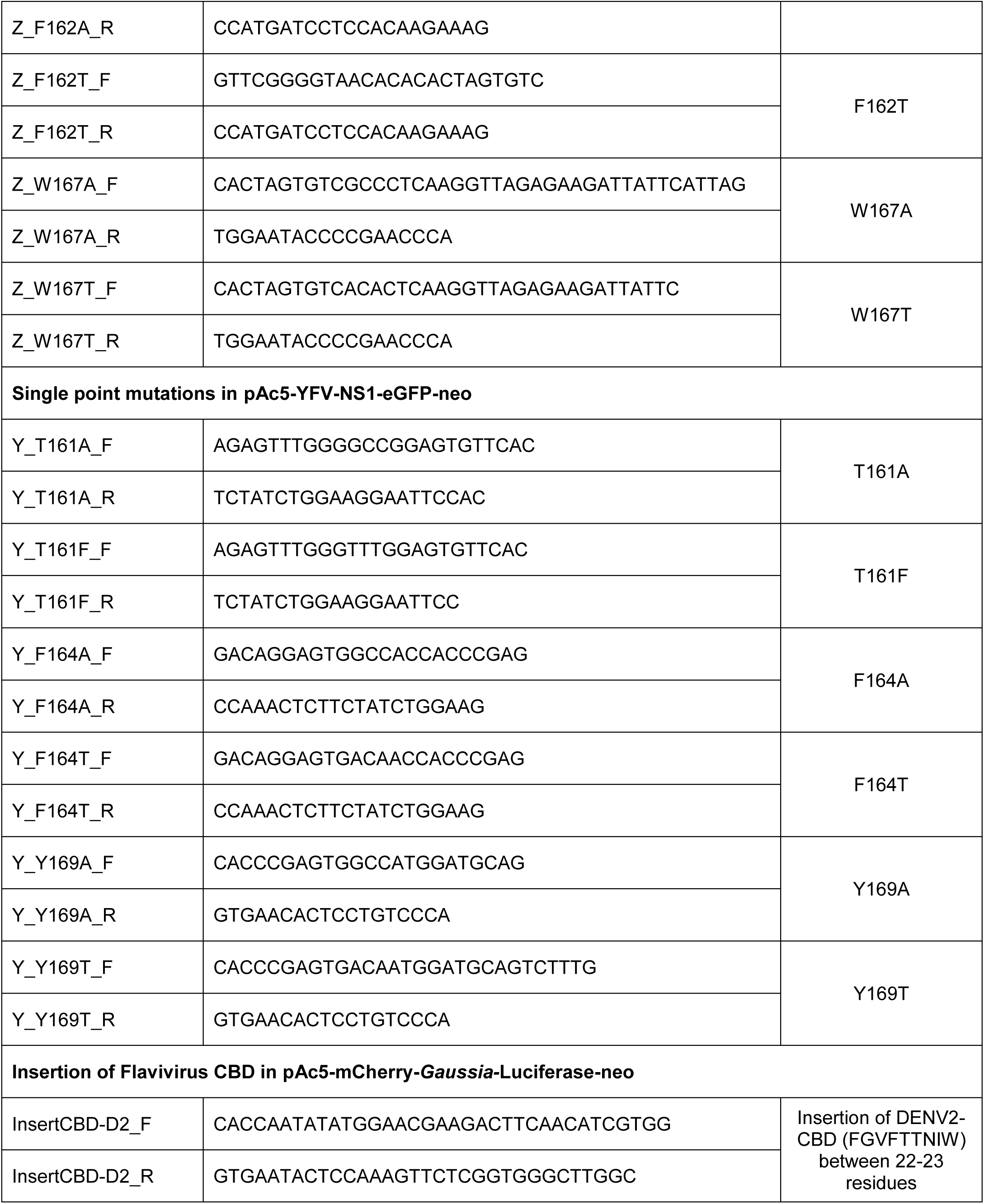
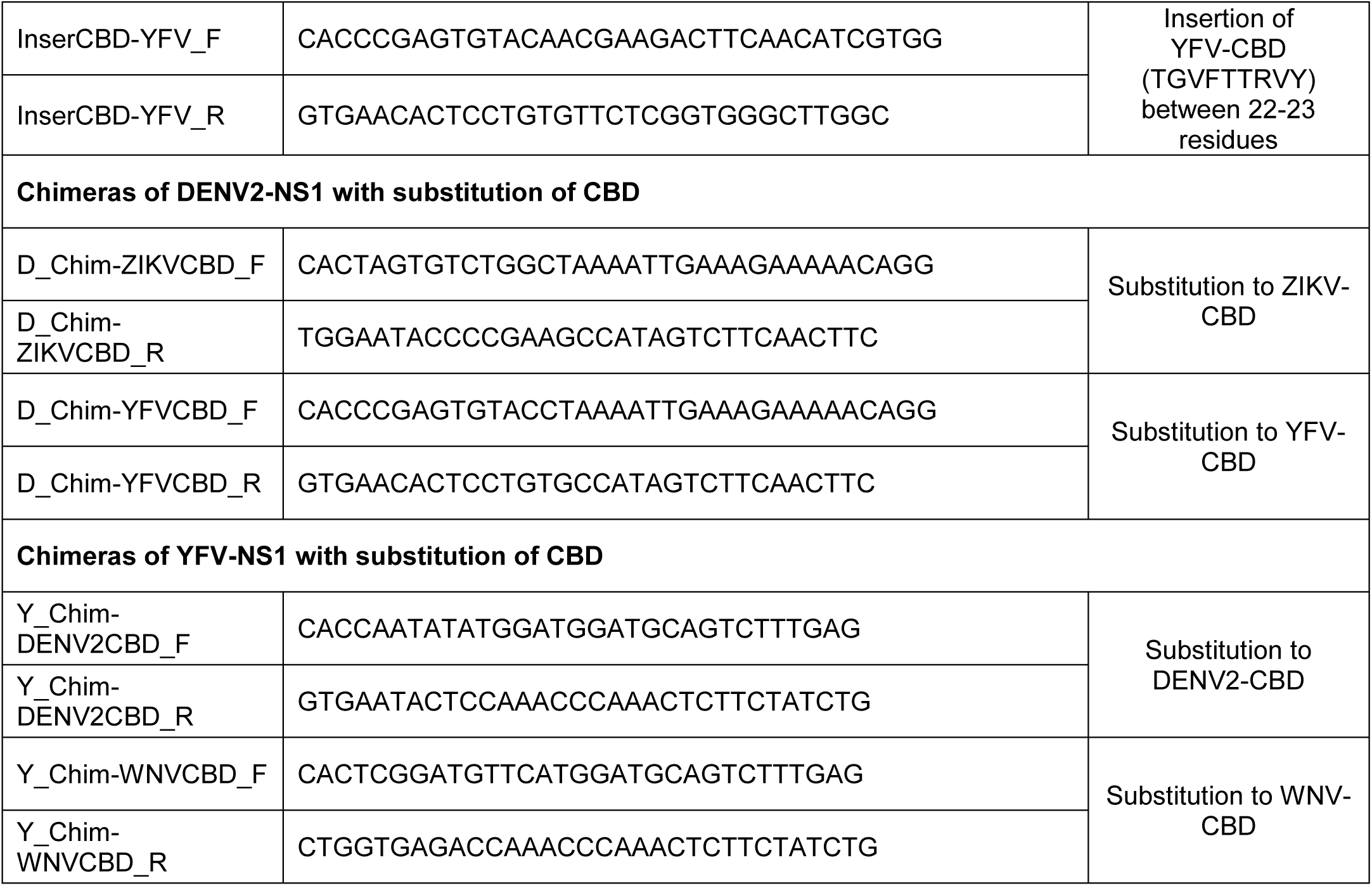
List of primers used for the site directed mutagenesis. DENV CBD located between 574-600 bp (188-196 aa); ZIKV CBD located between 1336-1362 bp (159-167 aa) and YFV CBD located between 493-519 bp (161-169 aa).

**Fig 1.**
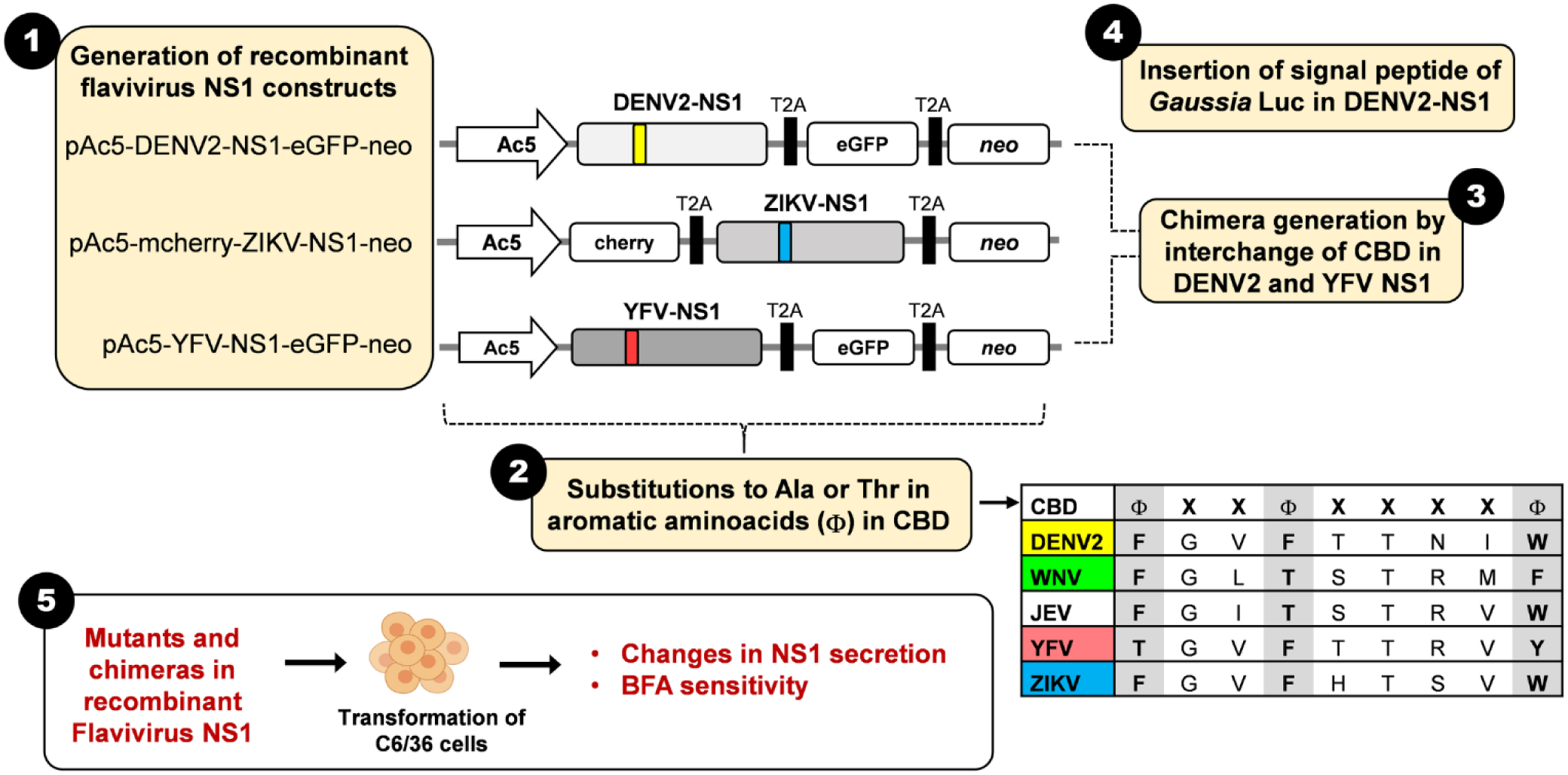
Schematic representation of the overall site directed mutagenesis strategy. **1. Generation of recombinant flavivirus NS1.** The details for the construction of recombinants NS1 are in Materials and Methods. Each construct was used as template for site directed mutagenesis. Actin5C promoter; cherry, red fluorescent protein; eGFP, eukaryotic green fluorescent protein; neo, resistance to neomycin, kanamycin and G418, each gene separated by a T2A peptide. Yellow and blue; DENV and ZIKV NS1 CBD sequences; red; YFV NS1 incomplete CDB. **2. Substitutions of aromatic residues in CBD to Ala or Thr.** CBD sequences of flaviviruses used in this manuscript. Note that WNV, and ZIKV present aromatic residues in all 3 positions. A few substitutions produce another flavivirus phenotypes in CBD sequence. **3. Chimera generation by interchange of CBD in DENV2 and YFV NS1.** The complete sequence of DENV2 CBD was substituted by the ZIKV or YFV CBD sequences. The complete sequence of YFV CBD was substituted by the DENV2 or WNV CBD sequences. **4. Insertion of signal peptide of *Gaussia* Luc in DENV2-NS1.** This insertion was introduced only in the DENV2-NS1 construct. **5. General experimental strategy with constructs.** All mutations, insertions and chimera constructions were transfected in C6/36 cells and NS1 secretion to the supernatants in cells treated or not with BFA determined by ELISA.

Aromatic positions in the DENV and ZIKV NS1 CBD were each change to Ala or Thr (Figure 1 and 2). Mutations to A were “non-sense”, but mutations F to T in the first aromatic position made the sequence YFV-like, and in the second aromatic position, WNV and JEV-like (Figure 1 and 2). In turn, the A to F mutation in the first aromatic residue of YFV NS1 result in the restoration of a full CBD. Mutations to both Ala or Thr in DENV NS1 resulted in a significant increase in the amount of secreted NS1. Interestingly, all three A and T DENV NS1 mutants showed increased secretion sensitivity to cell treatment with BFA, suggesting that the mutated DENV NS1 is at least partially secreted following a classical secretory pathway (Figure 2A). Results obtained with ZIKV NS1 mutants were less consistent; with increased NS1 secretion sensitivity to BFA cell treatment observed for only 3 of the 6 mutants constructed; that is mutants W167A and F159T and F162T, which convert them into a sequence similar to that of YFV and WNV (Figure 2B). Finally, in the case of the introduced mutations to YFV NS1, which sought to make secretion less sensitive BFA treatment, no significant changes in BFA sensitivity were observed; even the T161F mutation that converts the sequence into a complete CBD does not produce resistance to BFA (Figure 2C). In view of the results obtained with the single point mutations, complete substitutions of the DENV and YFV NS1 CBD were made (Figure 3). The DENV NS1 CBD was replaced for the complete YFV NS1 sequence, and the ZIKV NS1 CBD, as control. The YFV NS1 sequence was replaced by the DENV NS1 CBD and the WNV sequence, as control (Figure 3A). The results shown in Figure 3B indicate that the secretion of the DENV NS1 inserted with the YFV sequence, becomes sensitive to the treatment with BFA, while insensitivity to BFA is retained in the DENV NS1 with the ZIKV NS1 CBD. In addition, the insertion of the DENV NS1 CBD into the YFV NS1 sequence, renders the YFV NS1 secretion insensitive to BFA, while sensitivity for BFA treatment is still observed for the YFV NS1 inserted with the WNV sequence (Figure 3C).

**Figure 2.**
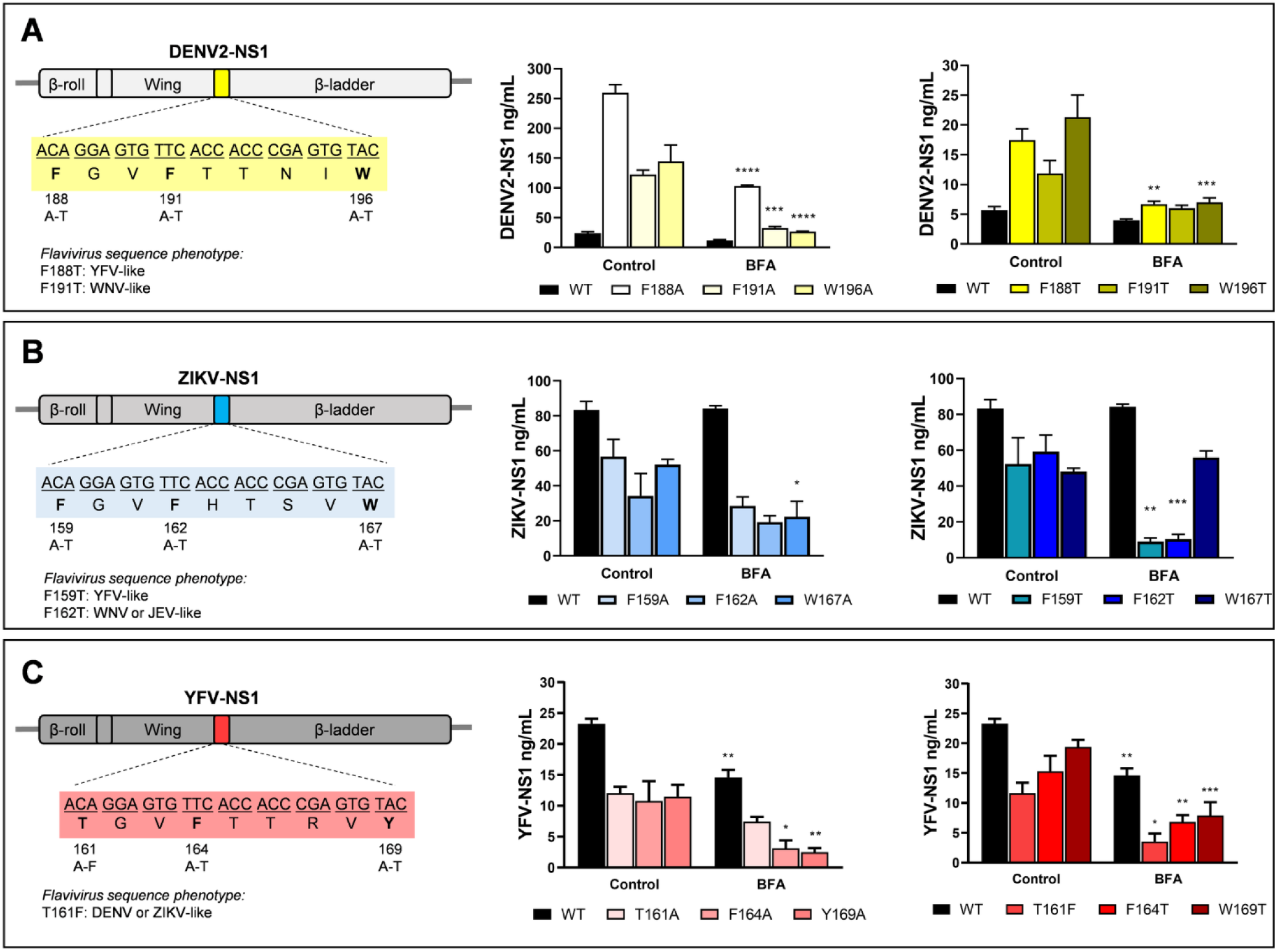
Secretory phenotype of recombinant DENV (A), ZIKV (B) and YFV (C) NS1 mutated in the CBD. *Left panels*, schematic representation of the mutations introduced in each of the NS1 genes. Secretion mutated NS1 to Ala (*central panels*) and Thr or Phe (*right panels*) in C6/36 cells treated or not with BFA. Twenty-four hours post-transfection, C6/36 cells were treated with DMSO (control) or with 25 µM BFA and the supernatants harvested after 48 h. Levels of secreted NS1 were measured by ELISA. Data are mean of 3 independent experiments ± standard error; significant differences between controls and BFA treatment are denoted by *(*p* < 0.0001).

**Figure 3.**
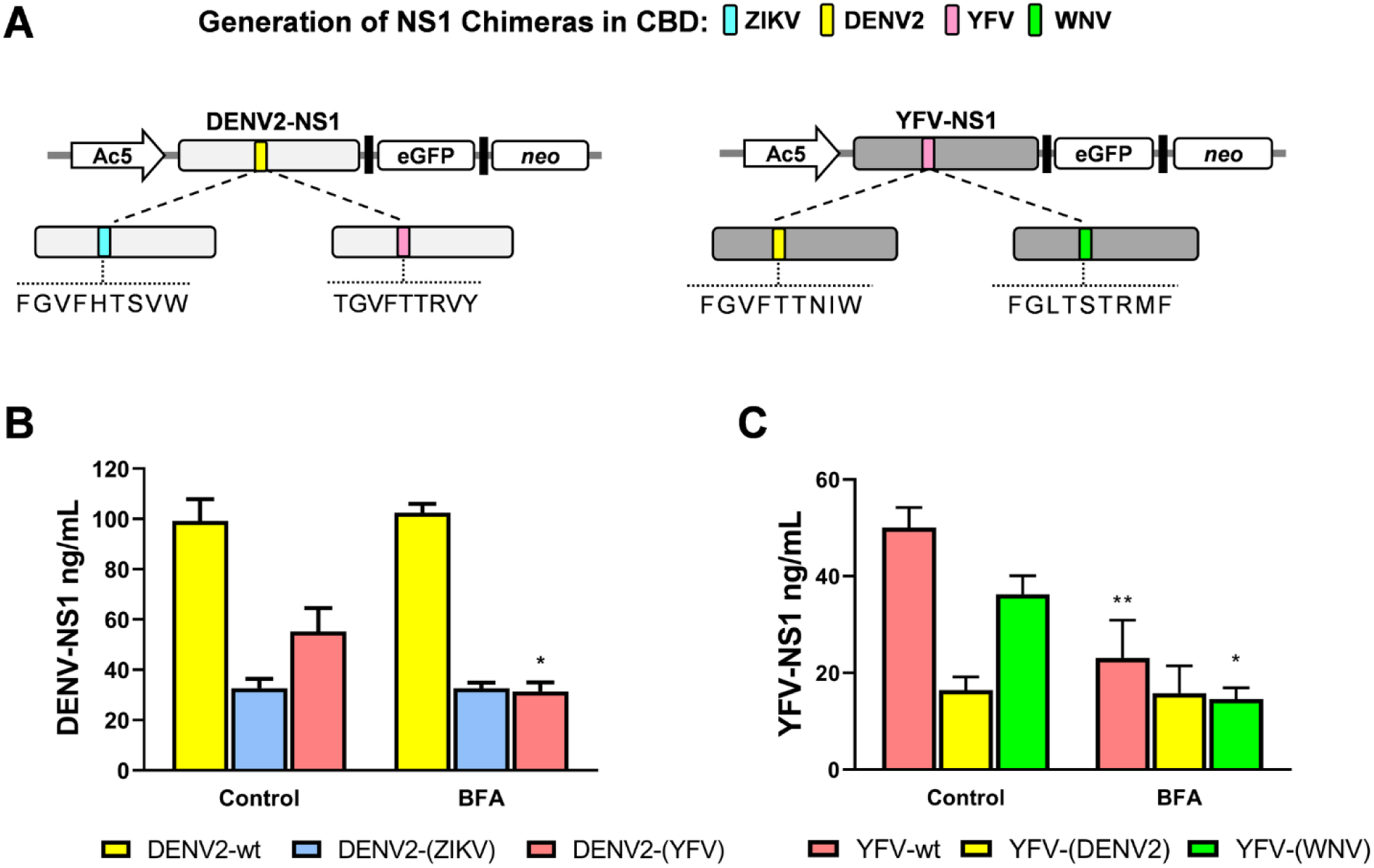
Secretory phenotype of CBD-chimeras of flavivirus NS1. **A.** Schematic representation of the chimeras generated from DENV (left) and YFV (right) NS1. Colors indicate the origin of the CBD. **B.** Secretion of DENV2-NS1 chimeras with ZIKV and YFV CBD. **C.** Secretion of YFV-NS1 chimeras with DENV and WNV CBD. Twenty-four hours post-transfection, C6/36 cells were treated with DMSO (control) or with 25 µM BFA and the supernatants harvested after 48 h. Levels of secreted NS1 were measured by ELISA. Data are mean of at least 3 independent experiments ± standard error; significant differences compared with controls are denoted by *(*p* < 0.0001).

In an attempt to force the secretion of DENV NS1 to follow a conventional secretory route, even in the presence of the CBD, the secretion sequence of *Gaussia* Luciferase, a protein that in mosquito cells is secreted by a conventional secretory route, was inserted at the N-terminal end of DENV NS1. These 17 amino acids (MGVKVLFALICIAVAEA) were inserted into the N-terminal end of NS1, which lacks a signal sequence secretion, by site directed mutagenesis (Figure 4A). However, as shown in Figure 4B, the insertion decreased the secretion of NS1 by more than 98%. Confocal microscopy analysis to determine where the GLuc-NS1 was being retained showed that the wild type protein was mostly located in the ER, while the (GLuc)-NS1 was observed not only in the ER but also in the Golgi-complex (Figure 4C and D). These results suggest that although the mutant protein reached the Golgi apparatus, it was retained there and not secreted, possibly due to retrograde transport to ER.

**Figure 4.**
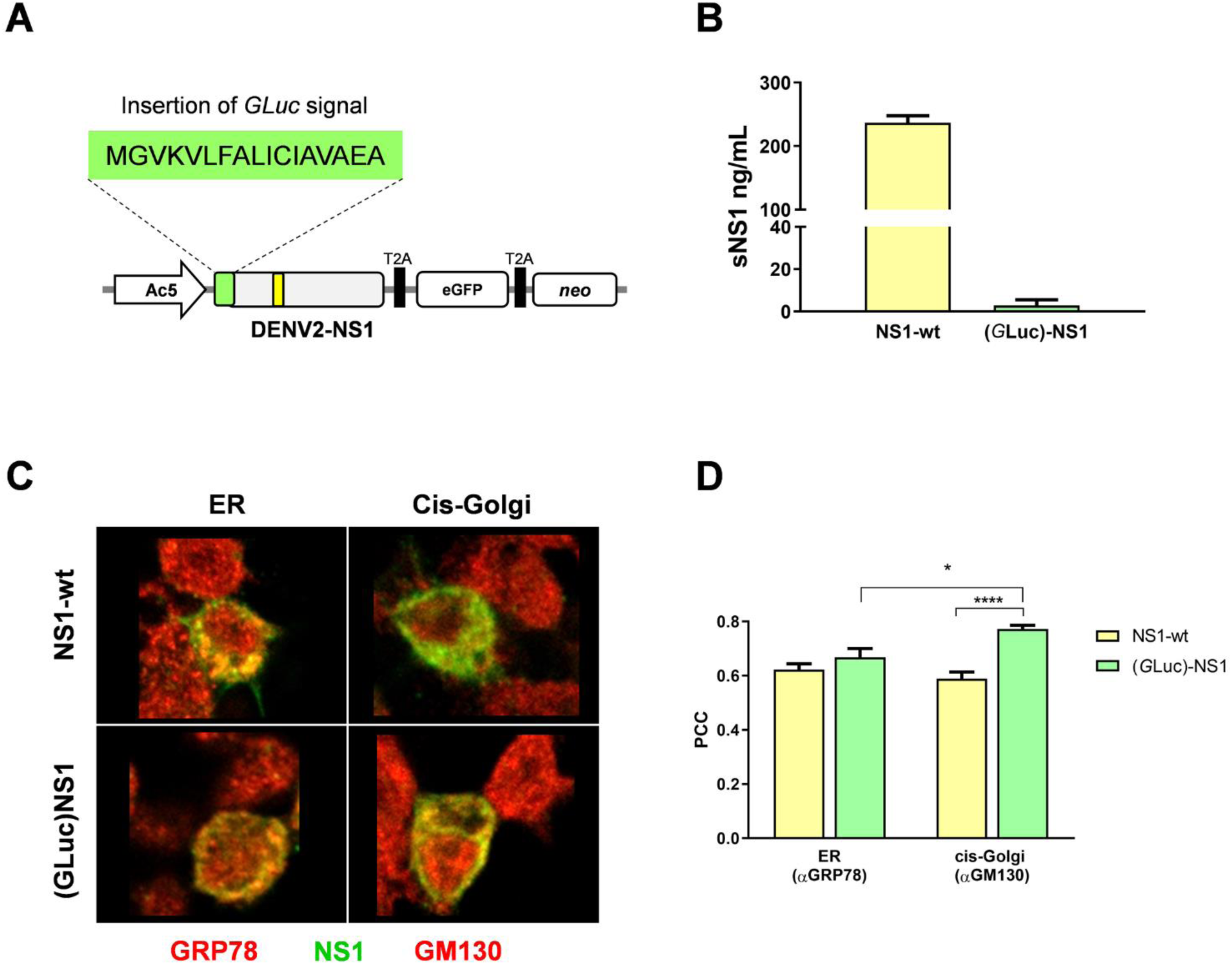
Secretory phenotype of DENV2-NS1 after addition of the signal peptide of *Gaussia* Luciferase. **A.** Schematic representation of the construction obtained after insertion of the *G*Luc signal peptide in the N-terminal of recombinant DENV2-NS1. The sequence MGVKVLFALICIAVAEA was introduced by site directed mutagenesis as described in the methods section. **B.** Secretion of GLuc-NS1 was measured in cells supernatants 48h post transfection. **C.** Co-localization between DENV2-NS1-wt or DENV2-GLuc-NS1 and ER (GRP78) and *cis*-Golgi (GM130) markers. Recombinants NS1 were transfected and 24hpt, cells were probed for NS1 (shown in green) and GRP78 or GM130 (shown in red). **D.** Pearson correlation coefficients (PCC) for organelle-NS1 were measured in at least 20 confocal independent images with 0.48 μM laser sections. The bars represent means ± standard error. Data was evaluated using the 2way ANOVA test and significant differences are denoted by *(*p* ≤0.05).

Finally, to evaluate the effect that the presence of the DENV CBD may have on a protein secreted by the classical route, a recombinant GLuc was constructed, into which the CBD of DENV NS1 was inserted (Figure 5A). Upon examination of the GLuc sequence, it was decided to insert the DENV CBD sequence, and that of YFV (as a negative control), between amino acids 22-23. This region is located after the signal peptide and before the catalytic domain of the luciferase and it is presumed to be exposed on the surface of the protein. Interestingly, the presence of DENV CBD made the GLuc secretion significantly less sensitive to BFA treatment; meanwhile the insertion of the equivalent YFV region did not produce any change (Figure 5C). These results again suggest that the presence of a functional CBD will adjust the protein secretory route, towards an unconventional route, in mosquito cells. Finally, co-localization experiments between GLuc and CAV-1 were performed to assess whether the presence of CBD would increase the interaction between Gluc and CAV-1. However, co-location analyzes, quantified by PCC, showed no evidence of a significant increase in the interaction between both proteins (Figure 5B).

**Figure 5.**
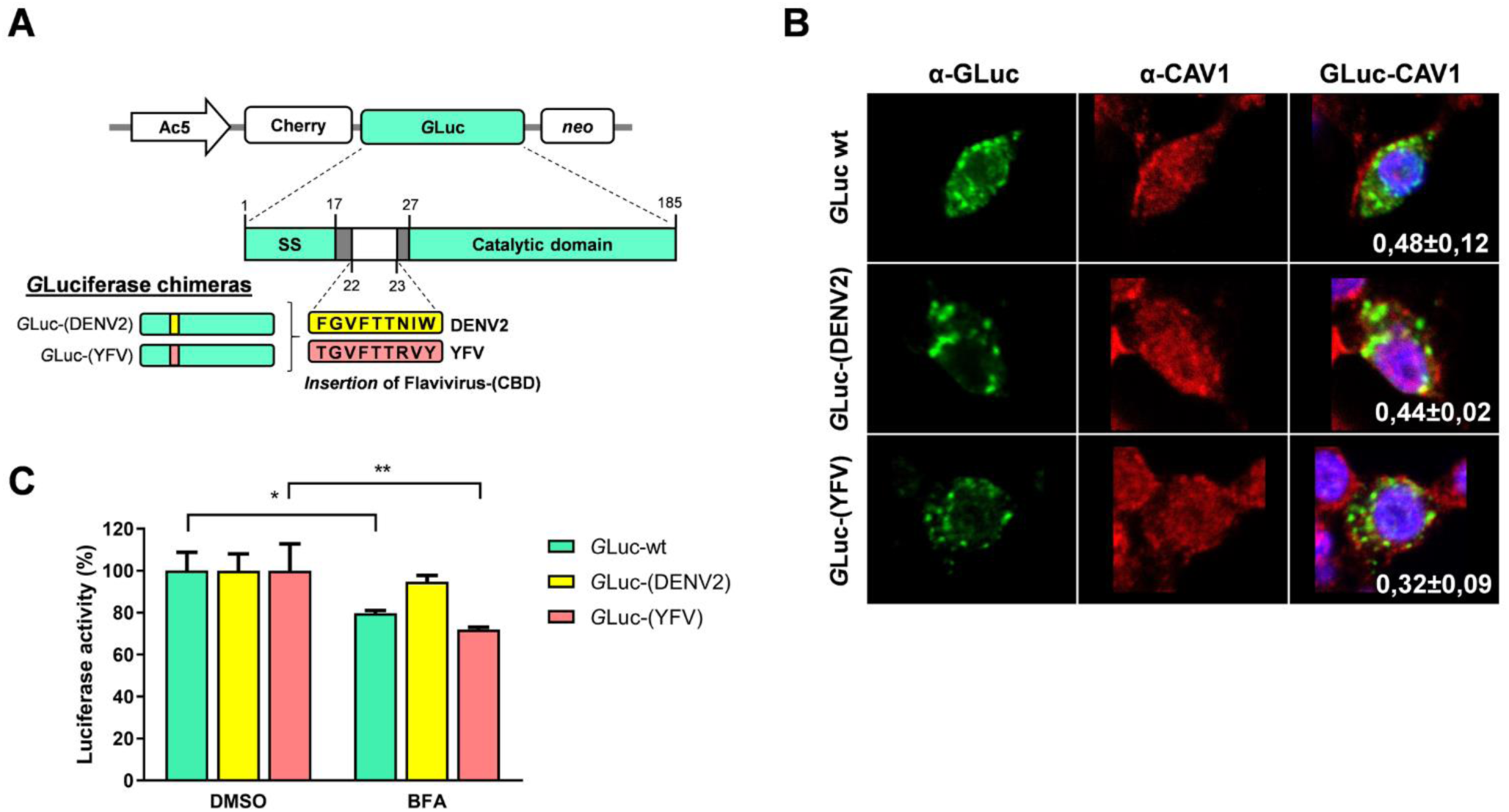
Secretory phenotype of *Gaussia* Luciferase after insertion of the DENV-NS1 CBD. **A.** Schematic representation of the construction of *G*Luc with the inserted DENV-NS1 CBD. Color boxes indicate the origin of the CBD. Ac5: Actin5C promoter; Cherry: red fluorescent protein; neo: geneticin resistance; SS: signal secretion, catalytic domain of luciferase. **B.** Confocal microscopy analysis of GLuc chimeras with CAV-1. Recombinant GLuc chimeras were transfected and 24hpt, cells were probed for *G*Luc (shown in green) and CAV-1(shown in red). The Pearson correlation coefficients (PCC) measured in at least 20 confocal independent images with 0.41 μM laser sections for each condition are shown. **C.** Secretion of *Gaussia* Lucifearse chimeras (DENV2 and YFV) in C6/36 cells treated for 48 hrs with BFA, or DMSO as control. Luciferase activity in control cells was taken as 100%. Experiments are based on at least three independent experiments with each chimera ± standard error; significant differences compared with controls are denoted by *(*p*< 0.0001).

## Discussion

The roles of NS1 in pathogenesis are associated to its presence in the extracellular and vascular space in the vertebrate host (9, 19, 20). However, the pathogenic effects within the mosquito vector are unknown. New findings have shown that flavivirus NS1 potently inhibits two important mosquito antiviral mechanisms (21). Although NS1 lacks a signal secretion in the N-terminal sequence, in vertebrate cells it follows a classical secretion pathway through Golgi to reach the extracellular space. In mosquito cells, an unconventional traffic route for secretion of DENV and ZIKV NS1 was observed, while in contrast, YFV NS1 seems to be secreted following the classical secretion route (11, 13). Given the presence of a conserved CBD in DENV and ZIKV NS1, but not in YFV NS1, a role for the CBD, as a molecular determinant, to direct the secretion fate within the *Aedes* endoplasmic reticulum architecture was proposed (13). Presumably, an active and exposed CBD in the hydrophobic region in the NS1 of flavivirus as DENV and ZIKV would facilitate the interaction with CAV-1 and directs the protein to an unconventional route in mosquito cells. In the present study, we employed a molecular genetic approach, using recombinant DENV, ZIK and YFV NS1s, to elucidate the role of the aromatic residues and the entire CBD in the secretory fate of flavivirus NS1 in mosquito cells.

By mutating any of the 3 aromatic amino acids of CBD in DENV2-NS1 that define the integrity of the CBD, the secretion of NS1 was changed from BFA cell treatment insensitive, to BFA sensitive, indicating that the mutants are now secreted, at least partially, following a classical secretory route through Golgi, as defined by BFA sensitivity (22–24). In addition, the DENV NS1 mutants, were secreted at significantly higher levels that the wild type protein. These changes in phenotype secretion suggest that the mutated DENV NS1 loses the ability to be recognized by CAV-1 or are recognized with less affinity, and therefore enter the route of classical secretion. Co-location analysis between point mutated DENV2 NS1 and CAV-1 showed no significant reduction in co-location levels (data not shown), a result which is compatible with only changes in affinity between NS1 and CAV-1. This observation is similar to a previous observations where the mutation of several residues within CBD of a cholesterol transport-protein, abrogated the export of cholesterol but did not change the binding of the protein to cholesterol (25).

However, the single point mutations within the Zika NS1, resulted in variable secretion phenotypes regarding BFA sensitivity. While mutations to Ala in the third aromatic residue and to Thr in the first and second aromatic residue in CBD of ZIKV NS1 induce a sensibility to BFA, other changes did not. The reasons for these differences with the DENV NS1 mutants are unknown, but differences in surface charges in the β-roll domain of the DENV and ZIKV NS1 have been reported (26), and despite the full conservation of the aromatic residues, 3 of the 4 amino acids found between the 2 and third aromatic residue are different, all of which may modulate the interaction between NS1 and CAV-1. In addition, the introduction of a T161F mutation to partially generate a CBD (aromatic amino acids in positions 1 and 2) into the YFV NS1, did not result in any change regarding BFA sensitivity, suggesting that the presence of a specific type of aromatic amino acid at the third position (like, Trp) and a complete CBD is required. All these results taken together suggest that single point mutations in DENV and ZIKV NS1 aromatic residues that disrupt the CBD result in changes in the traffic route of NS1 in mosquito cells.

Due to this behavior in the ZIKV and YFV NS1 single point mutants, we generated NS1 mutants where the complete CBD was exchanged. The DENV NS1 CBD was replaced by the YFV sequence (BFA sensitive) and the ZIKV CBD (BFA insensitive), as control; likewise, into the YFV NS1 the DENV CBD (BFA insensitive) was introduced and as a negative control, the sequence corresponding to WNV CBD. In both cases the presence of a conserved CBD, rendered the secretion of NS1 insensitive to BFA cell treatment, suggesting a secretion path change. Therefore, the presence of the two Phe, and the Trp at the end, appears to be necessary to direct the NS1 protein to follow an unconventional route. This conclusion was reinforced by the results obtained when the DENV CBD was introduced into the unrelated protein GLuc, which is secreted by the conventional secretory route. Surprisingly, when the DENV2 CBD was introduced into the recombinant GLuc, the secretion of GLuc gained insensitivity to BFA treatment, suggesting that it was now partially secreted by an unconventional secretion pathway. In the GLuc the CBD was introduced into an exposed area, and away the catalytic domain. This was an important consideration, since it has been described that the sole presence of a CBD, if not well exposed, does not guarantee that the protein will interact with CAV-1 (15, 27).

The results shown indicate that an active CBD is a molecular determinant guiding the secretion of DENV and ZIKV NS1 through an unconventional secretion pathway in mosquito cells. Yet, it is puzzling that the same sequence is not active in vertebrate cells, where the NS1 of these viruses is secreted via the ER-Golgi classical route (6, 12, 13). Unfortunately, the CAV-1 gene is *Aedes* mosquito have not been identify; however, docking simulations done with NS1 and CAV-1 from other insects, such as ticks (*Ixodes* sp., and *Sarcoptes* sp), showed a greater affinity of the DENV NS1 for these caveolins than for the human CAV-1 (data not shown). Thus, the mosquito cellular component acting as a sensor or recruiter for NS1 in could be the CAV-1 itself, although a role for cell architecture and other proteins, such as those of the *Sec* complex, cannot be discarded. The nearly complete abolition of secretion observed with the DENV NS1 modified with a secretion signal peptide, illustrates the complexity of the problem. Another interesting observation is that the CBD is fully conserved in DENV and ZIKV NS1, while incomplete in other flaviviruses such as YFV, JEV and WNV. Secretion of YFV NS1 have been observed in mosquito cells albeit a higher concentration was maintained as cell-associated rather than secreted into the extracellular milieu (28); moreover, no NS1 secretion is observed in WNV or JEV infected mosquito cells (29, 30). Thus, the presence of an active CBD and the recognition of NS1 by the mosquito caveolin may be crucial for NS1 secretion in the mosquito, and suggest unique roles for the DENV and ZIKV soluble NS1 in the mosquito (11, 21).

In summary, this work demonstrates that the sequence of PheXXPheXXXXTrp seems to play a role in determining that the unconventional secretory route of DENV and ZIKV NS1 in mosquito cells; resulting in interaction with CAV-1 and chaperones of the CCC. However, more research is needed to fully understand the viral and cellular factors that determine that the mosquito cell secretion machinery identify the CBD sequence and redirect the secretion pathway of NS1. Why these dramatic differences exist in the organization of the components of the secretion pathway between mosquitoes and in vertebrate cells is still an enigma. It is worth mentioning that the functions of the soluble NS1 in the mosquito are unknown and should be clarified; but may include the facilitation of the propagation of viral particles and the modulation of innate immunity. Finally, the manipulation of the lipid and cholesterol system in the mosquito can become a new target to reduce the secretion of NS1 and a new strategy to block the spread of mosquito-borne flaviviruses.

## MATERIALS AND METHODS

### Construction of recombinant flavivirus NS1 expression vectors and site directed mutaganesis

Ac5-STABLE2-neo was a gift from Rosa Barrio and James Sutherland (Addgene plasmid # 32426) (31). This plasmid was engineered to express recombinant flavivirus NS1 protein in mosquito cells. DENV2 NS1 (New Guinea C strain) gene was obtained from mammalian expression plasmid kindly donated by Dr. Ana Sesma (Icahn School of Medicine at Mount Sinai, New York). DENV2 NS1 was PCR-amplified using the following designed primers with directed cloning sites (underlined): *Forward primer-KpnI* (5’-GCTA<u>GGTACC</u>GCCACCATGGGATCACGCAGCACCTCACTGTCTGTG-3’) and *reverse-primer-NotI* (5’-CTTCGC<u>GCGGCCGC</u>GATCAGCTGTGACCAAGGAGTTGACCAAATTC-3’). DENV2-NS1 KpnI-NotI cassette was cloned into pAc5-STABLE2-Neo, generating the pAc5– (DENV2) NS1-GFP-Neo vector, under the promoter Actin5C (from *Drosophila melanogaster*) reported to be efficient in insect cells lines (31–33). The T2A peptide sequence derived from *Thosea asigna* (EGRGSLLTCGDVEENPGP) allowed multicistronic processing and the neomycin resistance gene (NeoR) confers resistance to G418 allowing stable mosquito cell lines.

The NS1 sequence from ZIKV (Mexican isolate, Asiatic linkage) and YFV NS1 (Brazilian yellow fever virus isolate) were synthesized *de novo* (GenScript, Piscataway, NJ). The synthetic ZIKV NS1 gene (GenBank accession number KY631493.1) was ligated as a XbaI/HindIII fragment into the similarly digested Ac5-stable2 expression cassette generating the pAc5–mCherry-(ZIKV)NS1-Neo vector. The synthetic YFV NS1 gene (GenBank accession number MH018093.1) was ligated as a KpnI/NotI fragment into the similarly digested Ac5-stable2 expression cassette generating the pAc5–(YFV)NS1-GFP-Neo vector.

The secretion signal from *Gaussia* Luciferase (GVKVLFALICIAVAEA) was inserted in the N-terminal sequence of DENV2-NS1 using designed primers between 15-16 nucleotide. Aromatic residues (Phe/Trp/Tyr) in the caveolin binding domain (CBD) within flavivirus NS1 plasmids were substituted to Ala or Thr with primers listed in Table 1To evaluate the effect of complete changes in the CBD in flavivirus NS1 sequence, we substituted the CBD from DENV2-NS1 to the CBD of ZIKV (TTCGGGGTATTCCACACTAGTGTCTGG), or the corresponding YFV sequence (ACAGGAGTGTTCACCACCCGAGTGTAC). These substitutions produced chimeras named DENV2-(ZIKV) and DENV2-(YFV), respectively. The CBD from YFV-NS1 construct was substituted to the CBD of DENV2 (TTTGGAGTATTCACCACCAATATATGG) or WNV (TTTGGTCTCACCAGCACTCGGATGTTC). These substitutions produced chimeras named YFV-(DENV2) and YFV-(WNV), respectively

All primers employed in substitutions, insertions and chimera construction were designed using NEBaseChanger v1.2.9 software and are listed in Table 1. Single point mutations, insertion of *G*Luc and chimera of CBD were introduced into the flavivirus NS1 constructs (DENV, ZIKV, YFV) using Q5® Site-Directed Mutagenesis Kit (NEB) used according manufacturer’s instructions.

### Insertion of flavivirus CBD in the Luciferase reporter

pAc5–mCherry-GLuc-Neo vector designed from a previous work was employed as a template for chimera construction by the insertion of caveolin binding domain of DENV2 and YFV (13). The selected region of insertion is located between the secretion signal domain (residues 1-17) and the luciferase catalytic domain (residues 28-185) (34). DENV2 CBD (FGVFTTNIW) encoded by sequence TTTGGAGTATTCACCACCAATATATGG, and YFV CBD (TGVFTTRVY) encoded by sequence ACAGGAGTGTTCACCACCCGAGTGTAC were introduced between the position 22 and 23 in the *G*Luc gene using primers designed using NEBaseChanger v1.2.9 software and primers are listed in Table 1. Insertion of CBD sequences were performed using the Q5® Site-Directed Mutagenesis Kit (NEB). Chimeras of *G*Luc were named *G*Luc-(DENV2) for DENV2 inserted sequence and *G*Luc-(YFV) for YFV inserted sequence.

Luciferase activity assay in the supernatants of transfected C6/36 cells with chimeras and wild type constructions were determined with Pierce™ Gaussia Luciferase Glow Assay Kit (Thermo Scientific). Percentage of luciferase activity was normalized with DMSO treatment.

### Cells and plasmid transfection

C6/36 cells from *Aedes albopictus* (ATCC^®^ CRL-1660™) were grown at 28 °C in Eagle’s Minimum Essential Medium (EMEM) (ATCC^®^ 30-2003™), supplemented with 5% fetal bovine serum (FBS) and 100 U/ml penicillin-streptomycin. Plasmid constructs were transfected into confluent monolayer of C6/36 cells using lipofectamine reagent Lipofectamine™2000 (Invitrogen). Each 24-well was transfected with 1 μg of plasmid DNA and 2 μL of Lipofectamine. After 5 h of transfection, cells were added EMEM with a final 10% FBS. After 24h, selective G418 was added to obtain stable C6/36 cell lines.

### Reagents and drug treatment

Brefeldin A (BFA) (B6542-Sigma-Aldrich) was dissolved in dimethyl sulfoxide (DMSO, ATCC^®^). BFA was used at a concentration of 7 μM in all experiments. Transfected cells were grown in 24-well plates and then, BFA was added to the cells in EMEM 5% FBS and G418 at 500 µg/mL. Incubation time was 48 hours or kinetic secretion assays at 28°C. After this time, cell supernatants were collected to measure secreted NS1. In other cases, cells were fixed and stained for immunofluorescence.

### Measurement of secreted NS1 protein

The presence of flavivirus NS1 in cell supernatants was measured using a non-commercial, in-house, ELISA. Briefly, ELISA 96-well plates (Nunc-Immuno™,Sigma-Aldrich®) were coated with 200 ng of purified anti-NS1 monoclonal antibody in carbonate buffer (0.05 M. pH 9.6) and incubated overnight at 4 °C. Non-specific binding was blocked by incubating with 100 μL/well of blocking buffer (PBS with 10% fetal bovine serum) for 1 h at 37 °C. At that point, supernatant samples were added (50 μL/well) and the plate incubated for 1 h at 37 °C. Then, 50 μL/well of anti-NS1 Mab (kindly donated by Eva Harris, Berkley University, CA) conjugated with biotin for 1 h at 37 °C were added, followed by 50 μL/well of streptavidin conjugated with HRP diluted 1:10.000 in PBS incubated for 1h at 37 °C. After each step, wells were washed by rinsing 3x with washing buffer (PBS with 0.01% Tween 20). The reaction was developed with the addition of 160 μL/well of TMB (Sigma-Aldrich®) for 15 min and stopped with the addition of 50 μL/well of 2M H2SO4. The color development reaction is proportional to amount of secreted NS1. The amounts of secreted NS1 was estimated in nanograms per mL using serial dilution of recombinant NS1.

### Confocal microscopy

Confluent cell monolayers, grown in 24-well plates containing glass coverslips, were transfected with vectors expressing flavivirus NS1 or *G*Luc. After the times indicated in the text, cells were fixed in paraformaldehyde 4% for 10 min. Cells were permeabilizated with 0.1% Triton X-100 for 10 minutes at room temperature and stained for DENV-NS1 using anti-NS1 Mab (kindly donated by Eva Harris, Berkley University, CA), anti-gaussia Luciferase (Pierce PA1181), anti-GRP78 (GTX22902 Genetex), anti-GM130 (G7295 Sigma-Aldrich), anti-CAV-1 (GTX89541 Genetex or sc-894 Santa Cruz), and Nuclei with DAPI. Anti-mouse Alexa-488 or Alexa-598, anti-goat Alexa-568 and Anti-rabbit Alexa-647 or Alexa-488 conjugated (Donkey pre-adsorbed, secondary antibodies, Abcam) were used at 1:800 dilution). Coverslips were mounted in Fluoroshield™ with DAPI (Sigma). Anti-GRP78 and anti-GM130 were used as endoplasmic reticulum and *cis*-Golgi markers, respectively. The images were analyzed using a LSM 700 confocal microscope. To evaluate the co-localization between proteins, Pearson correlation coefficients (PCC) were obtained from at least 20 confocal independent images (laser sections indicated in text) using the Icy image software and the co-localization studio plugin (35).

## Statistical analysis

Values of all assays were expressed as mean ± standard error of three independent experiments, each in triplicate or indicated in the text. Statistical analyzes were carried out using the GraphPad Prism version 6.01 software.

## Author contributions

Conceived and designed the experiments: RRR. JEL

Performed the experiments: RRR

Analyzed the data: RRR, JEL

Contributed reagents and materials: JEL

Wrote the paper: RRR, JEL

## Acknowledgments

This work was partially funded by CONACYT (Mexico) grant number CB-2015-1 254461 to JEL. Authors declare no conflict of interest.

